# Local differences in neuronal activation level decorrelate spatially coherent global fluctuations in gamma rhythms

**DOI:** 10.1101/150532

**Authors:** Kaushik J. Lakshminarasimhan, Nikos K. Logothetis, Georgios A. Keliris

## Abstract

Neuronal coherence is thought to constitute a unique substrate for information transmission distinct from firing rate. However, since the spatial scale of extracellular oscillations typically exceeds that of firing rates, it is unclear whether coherence complements or compromises the rate code. We examined responses in the macaque primary visual cortex and found that fluctuations in gamma-band (~40Hz) neuronal coherence correlated more with firing rate than oscillations in the local-field-potential (LFP). Although the spatial extent of LFP rhythms was broader, that of neuronal coherence was indistinguishable from firing rates. To identify the mechanism, we developed a statistical technique to isolate the rhythmic component of the spiking process and found that above results are explained by an activation-dependent increase in neuronal sensitivity to gamma-rhythmic input. Such adaptive changes in sensitivity to rhythmic inputs might constitute a fundamental homeostatic mechanism that prevents globally coherent inputs from undermining spatial resolution of the neural code.

---

Sensory neurons often exhibit changes in coherence in addition to firing rates, providing distinct substrates for representing information. Although the precise behavioral consequences of differences in the timescales of these two codes continue to be debated^1–3^, recent work suggests that they may operate in parallel to constitute a multiplexed temporal code^4^. A related but often overlooked issue is the compatibility of their spatial scales. Whereas spatially correlated firing is detrimental to the information capacity of rate codes, it is a defining aspect of synchrony-based codes. Entrainment of spikes from distinct columns can undermine the spatial resolution of the representation established by differences in firing rates.

Neuronal coherence in the gamma frequency range (30-90 Hz) is ubiquitous in the mammalian brain and has been implicated in a variety of functions including sensory processing^5–7^, attentional selection^8,9^, perceptual modulation^10,11^, working memory^12^, memory encoding and retrieval^13^ as well as neurological disorders like Schizophrenia and Parkinson’s disease^14,15^. Gamma oscillations are thought to be generated locally within the cortical microcircuit^16,17^ and have been reported to span hundreds of micrometers in the macaque brain. How does the spatial extent of gamma-band coherence compare to that of firing rates? What mechanisms, if any, help prevent global fluctuations in rhythms from compromising the integrity of the columnar organization?

To address these questions, we used multi-tetrode recordings to examine concurrent changes in spiking activity of single-units, local field potentials (LFP), and spike-field coherence (SFC) in the primary visual cortex of two rhesus macaques viewing monocularly presented gratings. Since the average synaptic activity is thought to be a major source of fluctuations in the LFP^18–20^, we expected the strength of LFP oscillations to best predict changes in the extent of synchronous firing estimated using SFC. In contrast, we found that gamma-band SFC was correlated more strongly with firing rate than strength of gamma rhythms in the LFP. We probed the underlying mechanism by partitioning the spiking process into two components – an asynchronous component that reflected stimulus-dependent changes in activation level, and a synchronous component driven by gamma oscillations in the LFP. We found that sensitivity of neurons to synchronous drive increased significantly with mean activity, and these changes specifically contributed to the stronger correlations between neuronal synchrony and firing rates. Such activity-dependent changes in sensitivity might constitute a fundamental mechanism that preserves the spatial resolution of the neural code by selectively entraining only highly activated neurons thus effectively decorrelating global, non-specific fluctuations in gamma rhythms.

## Results

### Single-unit spike-field coherence (SFC) in the gamma band

We recorded from 474 sites from two macaque monkeys viewing monocularly presented grating stimuli (**Methods M1 & M2**). A total of 811 single-units were isolated, among which ~75% (*n*=610/811) were deemed visually responsive (**Methods M3**). For each responsive unit, neuronal coherence was assessed by estimating spike-field coherences (SFC) between their spike trains and concurrently recorded field potentials (LFP) (**Methods M3 – Equation 1**). The response of a representative single-unit is shown in **Figure 1A**. The onset of the stimulus is accompanied by a sharp transient increase in the firing rate followed by a period of relatively sustained firing. As seen from the spike-field coherogram, there was a significant increase in coherence in the gamma range (30-45 Hz) following stimulus onset and this was most pronounced during the sustained period of neuronal firing (400-1000ms). Therefore, we confined all our analyses to this time window. Stimulus-evoked spiking activities of approximately 27% of visually responsive units (*n*=166/610; 119 from monkey D98; 47 from monkey F03) were found to exhibit significant gamma-band SFC (*p*<0.01; permutation test, **Methods M3**). SFCs were estimated between spike trains of the above single-units and LFPs measured from each of the simultaneously recorded sites (up to 6 sites in chronic; 4 sites in non-chronic recordings), yielding a total of 400 spike-LFP pairs. Since we want to study the fluctuations in gamma-band coherence, following results pertain to these 400 spike-LFP pairs unless stated otherwise.

**Figure 1.**
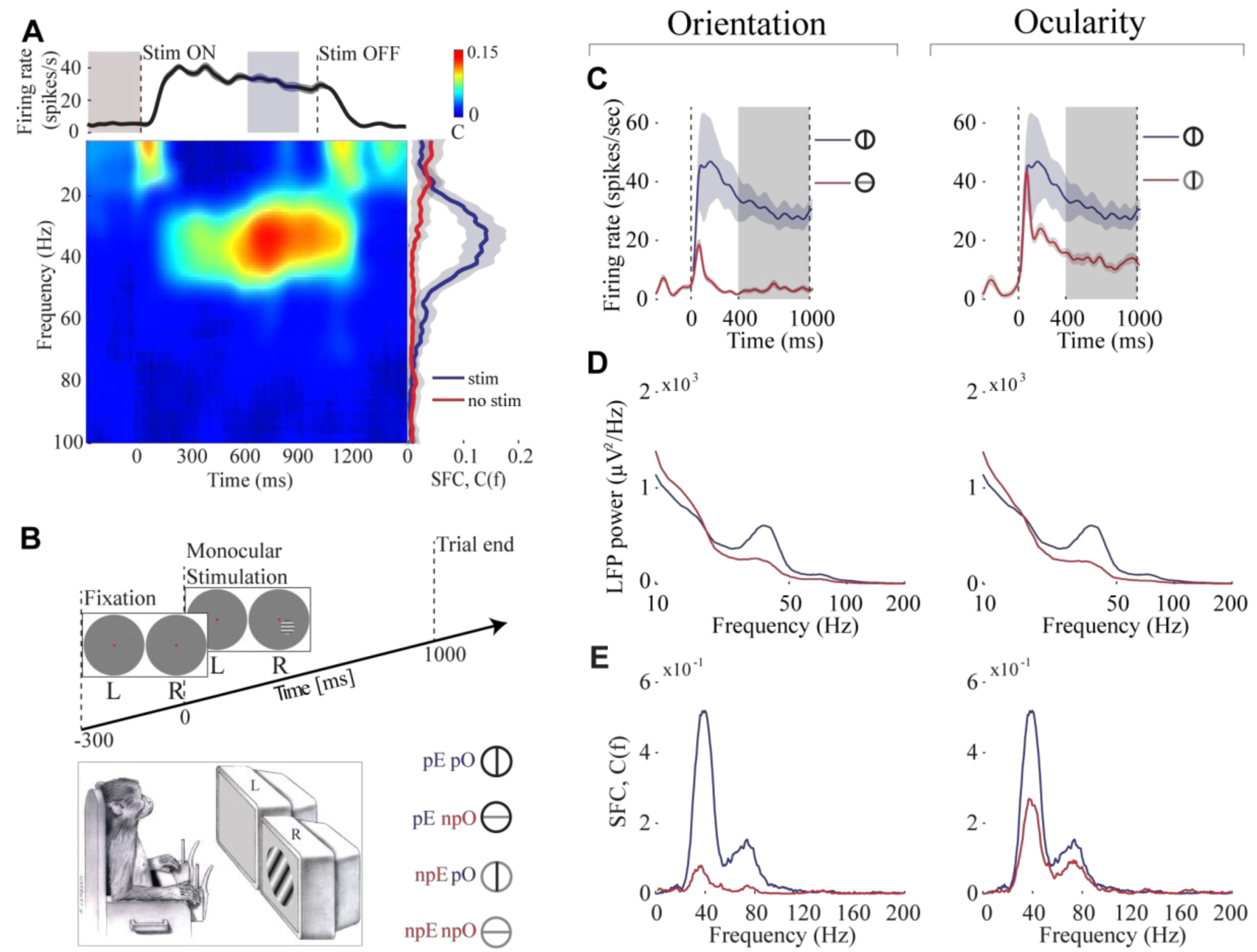
Modulation of neuronal coherence by stimulus preference. **(A)** Response of a representative spike-LFP pair. Top: Timecourse of trial-averaged firing rate, smoothed by Gaussian kernel with standard deviation of 25ms. Bottom left: Spike-field coherence (SFC) spectrogram during the concurrent period for frequencies up to 100 Hz, estimated by employing a 300ms long time window moved in steps of 1ms. Bottom right: SFC estimates before (red) and after (blue) onset of stimulus, computed using spikes and LFP during - 300 to 0ms (red region in top panel) and 600 to 900ms (blue region in top panel) windows respectively. Shaded areas depict 95% jackknife confidence intervals. (**B**) Experimental paradigm. Top: Each trial began with a central fixation dot presented to both eyes. After 300ms, a static circular sinusoidal grating of two possible orthogonal orientations was presented parafoveally to one of the eyes for a period of 1s. Bottom left: A cartoon illustration of the experimental set up. Stimuli were viewed dichoptically through a mirror stereoscope (not shown) that allowed the left (right) eye to view only the image on the left (right) display monitor labeled L (R). A calibration procedure carried out at the beginning of each session ensured that the displays were properly fused (**Methods M2**). Bottom right: Set of conditions employed in each experiment (for legend, see below). Different sessions employed different pairs of orthogonal orientations (not always vertical and horizontal) as determined by the preference of multiunit responses (**Methods M2**). (**C**) Left: Average firing rates of an example single-unit in response to gratings of preferred (blue) and nonpreferred (red) orientations presented to its preferred eye. The gray shaded region corresponds to the time window used for estimation of the LFP power and SFC. (**D**) Left: Power spectral density estimates (only 10-200 Hz shown for clarity) of a concurrently recorded LFP signal for the same pair of conditions. (**E**) Left: SFC between the single-unit spikes and the LFP signal. (**C-E -** Right panels) Similar quantities computed for the pair of conditions when a grating of preferred orientation was presented to either eye. (Legend: *pE* - preferred Eye, *npE* - nonpreferred Eye, *pO* - preferred Orientation, *npO* - nonpreferred Orientation, *pEpO* - preferred Eye preferred orientation, *pEnpO* - preferred Eye nonpreferred Orientation, *npEpO* - nonpreferred Eye preferred Orientation, *npEpO* - nonpreferred Eye nonpreferred Orientation).

### Stimulus-dependent changes in SFC are correlated with firing rates and LFP

We first analyzed stimulus-induced changes in SFC that accompany changes in firing rate and LFP for each spike-LFP pair by comparing responses under two sets of stimulus conditions (**Fig. 1B**): (1) the pair of conditions when either the preferred or nonpreferred orientation was presented to the neuron’s preferred eye (*pEpO* vs *pEnpO),* and (2) the pair of conditions when the preferred orientation was presented to either the preferred or nonpreferred eye separately (*pEpO* vs *npEpO).* These set of conditions were chosen to capture any effects that the differences in spatial scales between orientation and ocular columns might bear on our analyses.

We found that across the population of all spike-LFP pairs, changes in neuronal firing rate and gamma-band SFC were concomitant. Stimuli that elicited an increase in firing also produced higher SFC in most cases, i.e. degree of entrainment of spike trains to gamma-band LFP increased with spike density. On average, this was true both when neuronal response was manipulated by changing grating orientation as well as eye. As illustrated for one representative spike-LFP pair, increased spiking activity in response to the neuron’s preferred stimuli (**Fig. 1C**) is accompanied by an increase in LFP power around 40 Hz in a neighboring site (**Fig. 1D**). Moreover, this spike-LFP pair also exhibits an increased SFC in this frequency range (**Fig. 1E**).

Similar effects were observed in the majority of all analyzed spike-LFP pairs and are readily noticed in the population average of SFCs (**Fig. 2A, Supplementary Fig. 2A**), firing rates (**Supplementary Figures 1A,2B**), and LFP power spectra (**Supplementary Figures 1B,2C**). We tested the significance of stimulus-dependent changes in SFC (*C*) at the peak gamma frequency and firing rates (*R*) across the population of all spike-LFP pairs and found that both quantities increased significantly in response to the neuron’s preferred orientation as well as preferred eye (**Fig. 2B** – top panels; population rate: *R*_pEpO_=14.7±0.8 spikes s^-1^; *R*_pEnpO_=7.0±0.6 spikes s^-1^; *R*_npEpO_=9.1±0.7 spikes s^-1^, and population coherence: *C*_pEpO_=0.070±0.002; *C*_pEnpO_=0.013±0.001; *C*_npEpO_=0.039±0.002; p<10^-10^; Wilcoxon rank-sum test for the difference in medians between preferred and non-preferred responses). We examined the relationship between changes in firing rates and SFCs across the population, by comparing coherence modulation indices *M_C_* against corresponding rate modulation indices *M_R_* (**Methods M3 – Equation 2**). There was a strong positive correlation between *M_C_* and *M_R_* for the set of orientation (**Fig. 2B –** bottom left; Pearson’s correlation *r*=0.83; *p*<10^-10^) as well as ocularity conditions (**Fig. 2B –** bottom right, **Supplementary Figure 2D**; *r*=0.72; *p*<10^-10^). The above relationships were quantified using perpendicular regression and the slopes were found to be close to unity for the pair of orientation conditions (*M_c_*≈0.88*M_R_*; 95% confidence interval (CI): slope=[0.83 0.93]) as well as for ocularity (*M_c_*≈0.87*M_R_*; 95% CI: slope=[0.82 0.93]) suggesting that modulation in gamma-band synchrony and firing rate tend to be nearly identical.

**Figure 2.**
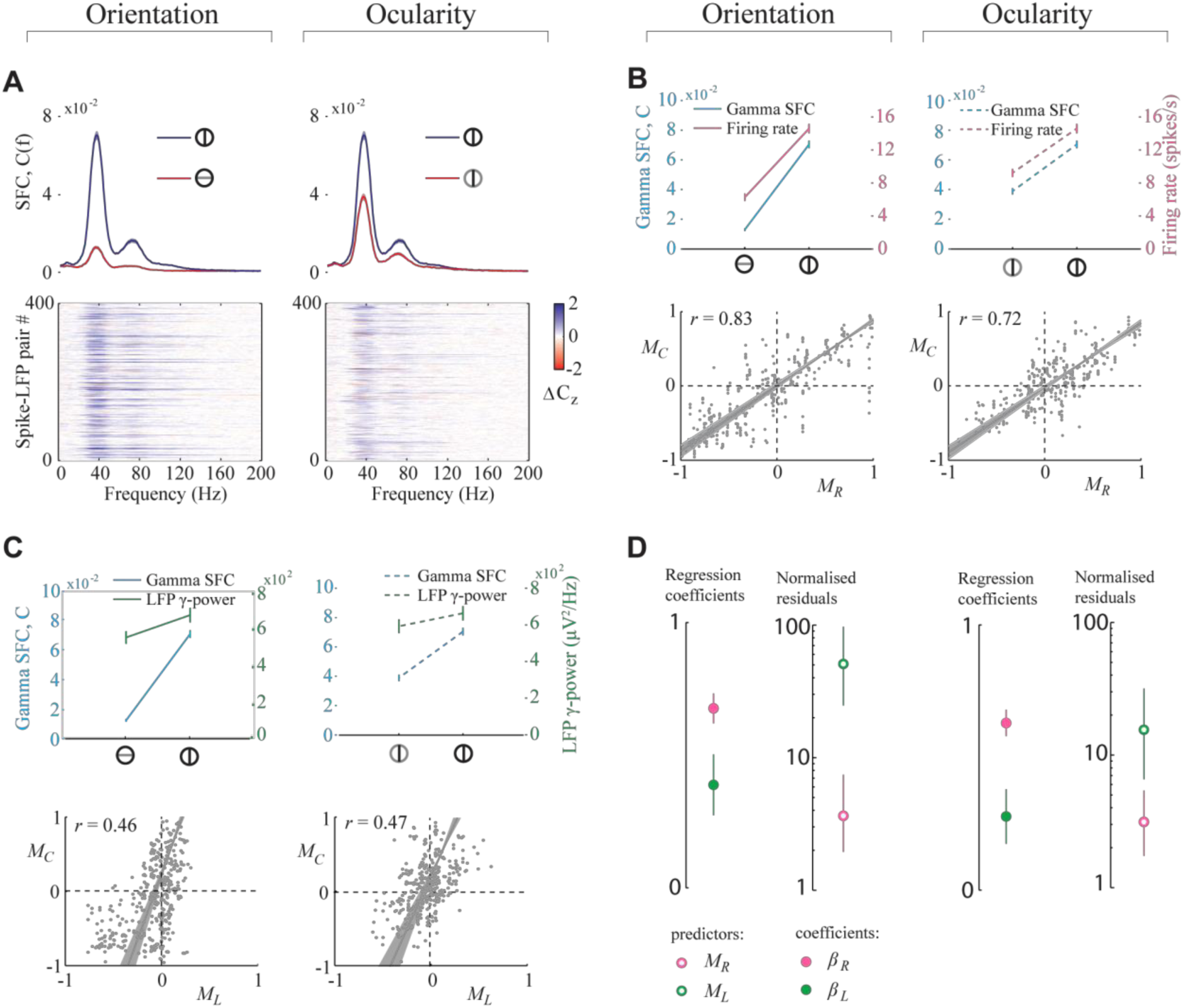
Stimulus-related changes in neuronal coherence are correlated with firing rate and LFP. **(A)** Top left: Average SFC across 400 spike-LFP pairs in response to the preferred (blue) and nonpreferred (red) orientations presented to the neurons’ preferred eye. Bottom left: Differences (preferred – nonpreferred) between the z-transformed SFCs of all individual spike-LFP pairs stacked vertically. Color bar indicates z-score differences (ΔC_z_) between the two conditions. (**B**) Top left: SFCs at the peak gamma frequency (cyan) in response to the pair of orthogonal orientation gratings – preferred (*pO*) and nonpreferred (*npO*) – shown to preferred eye (*pE),* averaged across 400 spike-LFP pairs. Firing rates (magenta) averaged across all single-units from those spike-LFP pairs, for the same pair of conditions. Error bars denote ± 1 standard error in mean (SEM). Bottom left: Rate modulation indices (*M_R_*) of orientation selectivity of the single-units from each spike-LFP pair (gray dot) plotted against the corresponding coherence modulation indices (*M_c_*). *r* denotes Pearson’s correlation coefficient between *M_c_* and *M_R_* and the solid gray line corresponds to the perpendicular regression between them. (**C**) Top left: Similar to (B), but showing concurrent changes in SFC (cyan) and gamma-band LFP power (green). Bottom left: Modulations in gamma-band LFP *M_L_* were significantly correlated with *M_c_*. (**D**) Left: The mean regression coefficients *β_R_* (magenta) and *β_L_* (green) for predictors *M_R_* and *M_L_* respectively, obtained by multiple linear regression. Error bars denote bootstrapped estimates of standard deviation. Plot on the right shows normalized residuals (**Methods**) resulting from regressing *M_c_* separately against either *M_R_* (magenta) or *M_L_* (green) shown as median ± central-quartile range. (**A**-**D** - Right panels) Similar quantities computed for the pair of conditions when a grating of preferred orientation was shown to either eye.

Next, we examined whether changes in SFC are related to changes in the strength of gamma-band LFP. If LFP primarily reflects the average synaptic input to neurons in the local circuit, then an increase in gamma-band power of the LFP should predict an increase in the extent of coherent firing of those neurons. Indeed, stimulus-related modulations of SFC were largely congruent (**Fig. 2C** - top panels) and significantly correlated with those of the LFP gamma-band power *M_L_* in both pairs of stimulus conditions (**Fig. 2C** – bottom panels, **Supplementary Figure 2E**; orientation: *r*=0.46; *p*<10^-10^; ocularity: *r*=0.47, *p*<10^-10^). Surprisingly however, these correlations were significantly smaller in magnitude than their correlation with firing rate (*p*<10^-10^, one-tailed two-sample *t*-test for the difference in correlations). Since majority of spike-LFP pairs were comprised of spikes and LFPs recorded at different sites, we wanted to know whether weaker correlation of neuronal coherence with LFP was due to distance effects. We tested this by grouping spike-LFP pairs based on the electrode separation between singleunit spikes and LFPs. We found that the correlation coefficients were not significantly different across groups thus ruling out this possibility (**Supplementary Fig. 3**).

To further quantify the differences between the effect of firing rate and strength of gamma-band LFP on neuronal coherence, we performed a multiple linear regression of *M_c_* with both *M_R_* and *M_L_* as simultaneous predictors (**Methods M3 – Equation 3**). We found that the regression coefficients on *M_R_* were significantly larger than on *M_L_* (**Fig. 2D**; Orientation: *β_R_*=0.68±0.06, *β_L_*=0.39±0.1, *p*<10^−10^, Ocularity: *β_R_*=0.63±0.05, *β_L_*=0.29±0.1, *p*<10^-10^; one-tailed *t*-test comparing regression coefficients *β_R_* vs *β_L_*), indicating that changes in firing rate rather than LFP, better explained the stimulus-related variability in gamma-band coherence. In fact, changes in SFC predicted by a linear regression model solely with *M_R_* as predictor generated residuals that were about five to ten times smaller (*p*<10^-5^, *t*-test) than those predicted using *M_L_* alone (**Fig. 2D**). Nevertheless, regression coefficients of the two predictors were significantly above zero implying that firing rate and the strength of LFP oscillations both carried independent information about changes in neuronal coherence.

Above results demonstrate that changes in SFC are correlated both with firing rate and gamma-band LFP, when those changes were induced by stimulus manipulation (either the orientation or the eye). To test whether these quantities are also correlated in the absence of stimulus change, we computed their correlated variability across trials within each stimulus condition. To do this, we estimated the correlation between trial-by-trial pseudo-SFC (pSFC) values (**Methods M3 – Equation 4**) and concomitant fluctuations in firing rate (*ρ_RC_*) as well as gamma-band LFP power (*ρ_LC_*) at all frequencies. If correlations were exclusively due to stimulus change, one would expect them to vanish when the measurements are conditioned on the stimulus. In contrast, across the population of all spike-LFP pairs, both *ρ_RC_* and *ρ_LC_* were found to be significantly positive in the gamma-band (*p*<10^-3^, Fisher’s combined probability test) within each stimulus condition (**Supplementary Fig. 4A, B**). Thus correlations persisted in the absence of stimulus change. Furthermore, the strength of correlation at the peak gamma frequency was found to depend on stimulus identity such that stronger input drive elicited greater correlations (*pEpO* > *npEpO* > *pEnpO*; *p*<10^-5^, Kruskal-Wallis test for correlations vs stimulus). Specifically, presentation of the neurons’ preferred stimuli increased the correlation between neuronal synchrony and firing rate (**Supplementary Fig. 4A** – inset; *ρ_RC_* : *pEpO* - 0.16±0.01, *npEpO* - 0.13±0.01, *pEnpO* - 0.11±0.01), as well as the correlation between synchrony and gamma-band LFP (**Supplementary Fig. 4B** – inset; *ρ_LC_* : *pEpO* - 0.15±0.01, *npEpO* - 0.10±0.01, *pEnpO* - 0.04±0.01). These results have two key implications. First, correlation between firing rate, LFP and neuronal coherence is not simply due to these quantities all being identically tuned to stimulus, but is likely a signature of an intrinsic mechanism that couples changes in firing rate and gamma-band LFP to neuronal coherence. Second, this mechanism might be sensitive to stimulus drive such that a stronger drive leads to a tighter relationship between these measures.

### Neuronal coherence and LFP are spatially dissociated

Why do firing rates, rather than LFP rhythms, better predict changes in neuronal coherence? We have previously shown that, for stimuli presented within the classical receptive field, LFP reflects activity spanning the order of ocular dominance columns^21^. Since coherence was more correlated with firing rate, we hypothesized that changes in the global strength of rhythmic synaptic input reflected in the LFP must be gated by mechanisms of a spatially local origin to ultimately limit the spatial extent of synchronous firing. If this is true, then the spatial scale of neuronal synchrony would be determined primarily by that of the firing rate code, and would hence be much smaller than the scale of LFP rhythms.

To test this, we estimated the pairwise correlation between SFCs of simultaneously recorded neurons as a function of the distance between electrodes from which the neurons were recorded (**Fig. 3**, cyan). Likewise, we also estimated correlated variability in firing rates of all pairs of neurons (**Fig. 3**, magenta) as well as correlations in gamma-band LFP powers at the respective electrode sites from which those neuronal pairs were recorded (**Fig. 3**, green). In each case, we computed both stimulus-induced (also called ‘signal correlation’, **Fig. 3A**) and stimulus-independent (‘noise correlation’, **Fig. 3B**) components of the correlation. To assess the effect of spatial separation, we analysed how correlations in each of the three measures (gamma LFP, firing rate, and gamma SFC) changed with distance by dividing the pairs of electrodes into two groups of roughly equal size: those that were nearby (<=200 μm) or far away (>200 μm).

**Figure 3.**
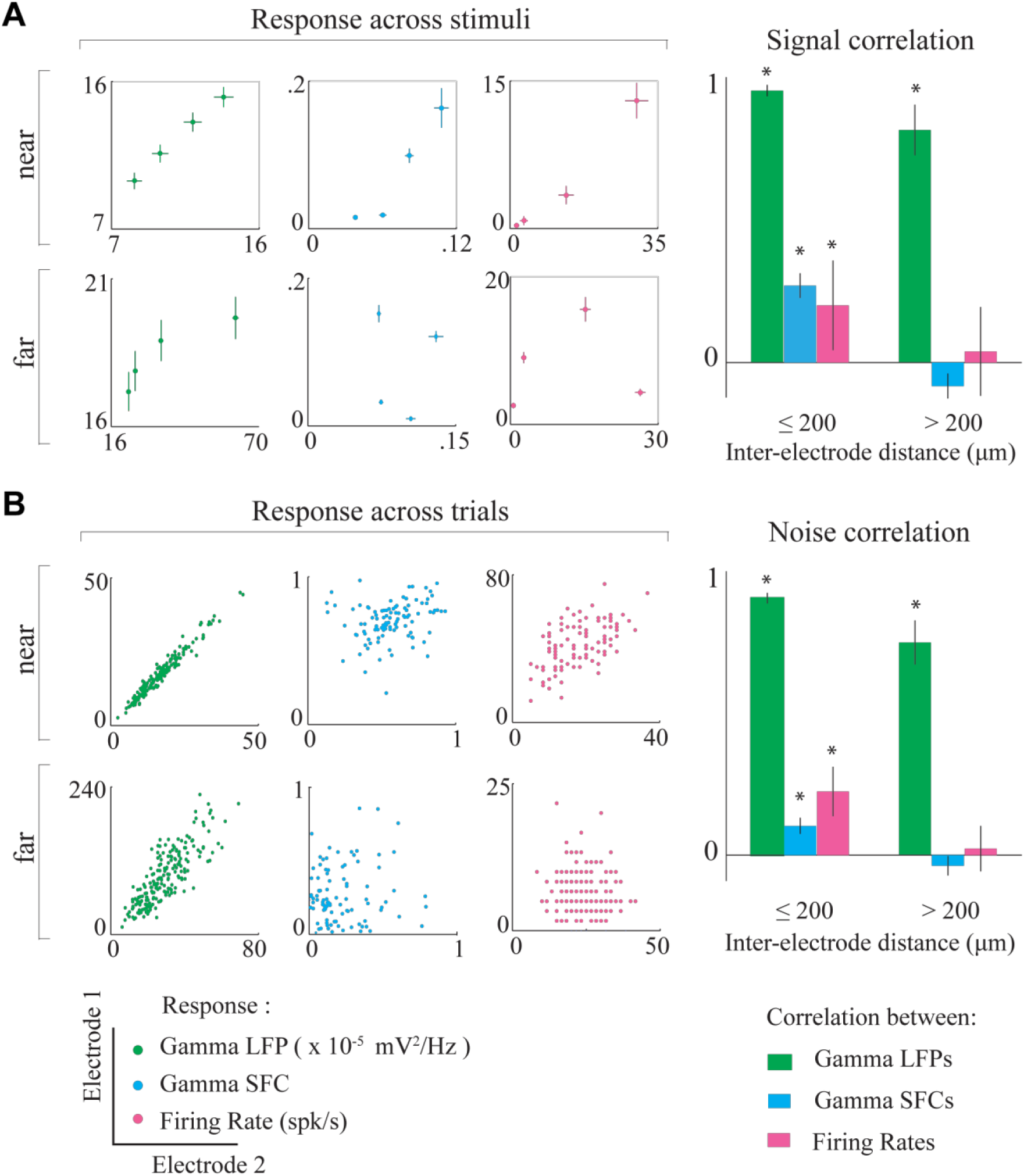
Spatial scales of LFP and coherence. **(A)** Left: Trial-averaged responses to four different stimuli (see Fig. 1B) recorded at two pairs of example sites that were nearby (top panels) or far away (bottom panels). Whereas gamma-band LFP power, gamma-band SFC, and firing rates were all similarly tuned across nearby sites, only gamma-band LFP was similar across more distant sites. Right: Average signal correlations between pairs of nearby (<200 pm) and distant (>200 pm) sites for each of the three response measures. Whereas the tuning of gamma-band LFP remained significantly correlated between distant sites, those of SFC and firing rate were not significantly different from zero for distant sites. **(B)** Left: Trial-by-trial responses to one of the stimuli at two pairs of example sites. Unlike LFP oscillations, fluctuations in SFC and firing rates were uncorrelated at the pair of distant sites. Right: Average noise correlations showed similar distant-dependent effects as signal correlations. Error bars denote ±1 SEM (^*^ *p*<0.01, two-sided sign test for median correlation of zero).

The panel on the left in **Figure 3A** shows concurrent stimulus-induced changes in gamma-band LFP (green), SFC (cyan), and firing rate (magenta) at pairs of nearby (top) and far away (bottom) electrode sites. Whereas signal correlations between gamma-band LFPs were high at both pairs of locations, correlations in SFC and firing rates were both only significant between neurons in nearby sites. This trend was observed across our dataset (**Fig. 3A** - right). The similarity in tuning of gamma-band LFP power remained large and significantly above zero across long distances (nearby pairs: *r*=0.95±0.04, *p*<10^-10^, two-sided sign test; distant pairs: *r*=0.81±0.18, *p*<10^-10^), whereas (signal correlations in both SFCs (nearby pairs: *r*=0.27±0.06, *p*<10^-10^; distant pairs: *r*=-0.08±0.06, *p*=0.053) and firing rates (nearby pairs: *r*=0.2±0.15, *p*=0.0015; distant pairs: *r*=0.04±0.16, *p*=0.23) were only significant between neuronal pairs in nearby sites. The magnitude of noise correlations exhibited a similar trend (**Fig. 3B**). Significant correlations were found between trial-by-trial changes in gamma LFP regardless of distance (nearby pairs: *r*=0.90±0.03; distant pairs: *r*=0.75±0.16). On the other hand, SFCs (nearby pairs: *r*=0.10±0.05; distant pairs: *r*=-0.04±0.04) and firing rates (nearby pairs: *r*=0.22±0.08; distant pairs: *r*=0.02±0.07) of neurons at distant sites were both uncorrelated.

### LFP rhythms are robustly correlated with membrane potential

Above results suggest that although neurons with large separation likely receive similar rhythmic synaptic inputs as implied by the long-range correlations in the LFP, their outputs are incoherent. Instead the spatial scale of neuronal coherence is similar to that of firing rates. This raises the possibility that globally correlated rhythmic synaptic inputs are gated by spatially local mechanisms that are sensitive to neurons’ activation level, to ultimately restrict the extent of rhythmic synchronization of their outputs.

However a alternative explanation for dissociation in the spatial scales of neuronal coherence and LFP is that gamma activity in LFP does not reflect the strength of rhythmic synaptic inputs to the neurons in our recording. If rhythmic input to individual neurons was more local than the LFP suggests, then the lack of a strong relation between neuronal synchrony with LFP could be explained away without the need to invoke any local mechanism. A direct way to test this alternative would be to compare LFP rhythms against oscillations in membrane potential of the neurons in our dataset. Such a direct comparison was not possible due to difficulties associated with preforming stable intracellular recordings in awake macaques. Instead, we indirectly estimated the correlation between membrane potential and LFP by computing spike-triggered average (STA) of the LFP. We computed normalised STAs for each neuron under all stimulus conditions by dividing the STA by the standard deviation of the LFP under the corresponding condition (Methods M3). A previous study involving simultaneous intra- and extracellular recordings has shown that this normalised STA is essentially equal to the cross-correlation between the neuron’s membrane potential and the LFP^22^. Therefore we used the amplitude of the normalised STAs to assess whether gamma-band LFP is a good predictor of gamma rhythmic synaptic inputs to the individual neurons in our dataset. Figure 4A shows normalised LFP STAs estimated using spikes of one example neuron under the pair of orthogonal stimulus conditions. STAs of this neuron revealed a substantial correlation between the LFP and synaptic inputs in the gamma range, and the magnitude of this correlation was similar across stimulus conditions. This was true on average across the population of all neurons (STA amplitudes: *pEpO* - 0.44±0.28, *npEpO* - 0.44±0.26, *pEnpO* - 0.42±0.27). The median amplitudes of STA were not significantly different across orientations of the grating (**Fig. 4B**; *p*=0.50, Wilcoxon rank-sum test) or the eyes that were stimulated (**Fig. 4B** – *right*; *p*=0.57). If gamma rhythmic input to individual neurons was already localized in space and varied concomitantly with firing rates, then its correlation with the global LFP signal should change with stimulus. In contrast, our results show that gamma-band LFP can predict the strength of rhythmic synaptic input equally well under all stimulus conditions. This robustness suggests that rhythmic input to neurons must be broadly correlated across space and argues for an active mechanism that decorrelates the neuronal outputs.

**Figure 4.**
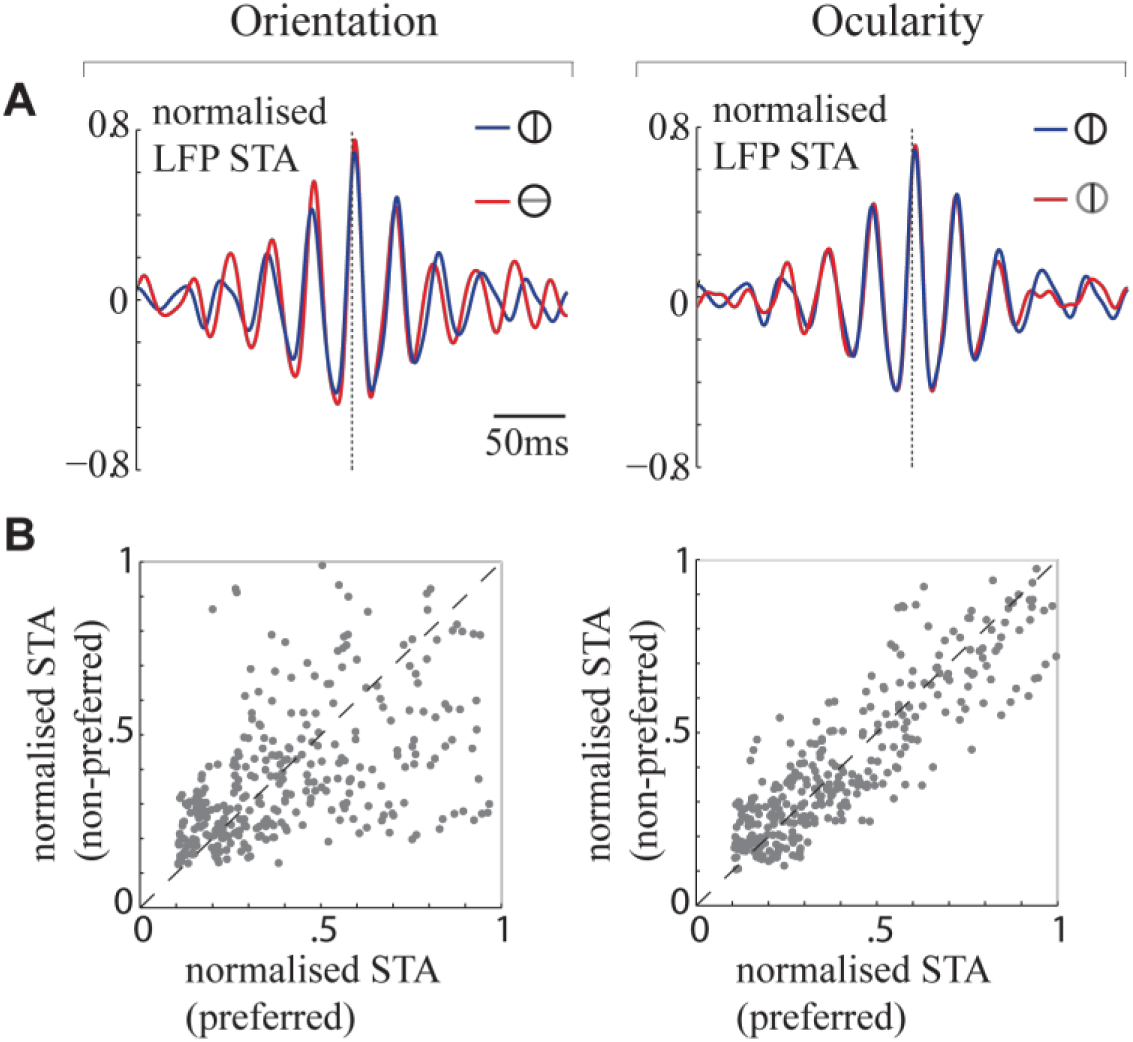
Comparison of the spike- triggered average (STA) of the LFP across stimulus conditions. (**A**) *Left*: Average LFP waveforms (normalised by the standard deviation of the LFP) within a 300ms window centered on the time of spikes (dashed vertical line) emitted by an example neuron in response to the presentation of its preferred (blue) and nonpreferred (red) orientations. *Right:* Waveforms computed around spikes emitted in response to stimulus presented to the neuron’s preferred (blue) and non-preferred (red) eye. (**B**) Comparison of the amplitude of the normalised LFP STAs across the pair of orientation (left) and ocularity (right) conditions for the population of all neurons. STA amplitudes are not systematically modulated by neurons’ stimulus.

### Activity-dependent increase in sensitivity to synchronous input

In order to account for our experimental findings, we considered a simple linear model in which the spiking process **r** is given by **r** = **A** + *g* · **ψ** where **A** denotes asynchronous activation with mean *a* that reflects the net excitatory synaptic drive, **ψ** is the rhythmic synaptic input, and the gating parameter *g* represents the sensitivity to synchronous input. The activation parameter *a* determines the average firing rate of the neuron, while *g* determines how well synchronous input **ψ** is encoded in the temporal pattern of spikes. In this model, rhythmic input drives coherence in the output while asynchronous activation reduces it. Thus neuronal coherence should be positively correlated with the strength of rhythmic input but anti-correlated with the overall activation level reflected in the firing rate (**Supplementary notes, Supplementary Fig. 5A**), a prediction at odds with our experimental results. However, if there is an activity-dependent increase in sensitivity to synchronous input (*g* ∝ *a*) that facilitates the transfer of synchrony, synchrony at the output could increase with firing rate (**Supplementary notes, Supplementary Fig. 5B**). Therefore, we wanted to know whether stimulus-related increase in neuronal coherence in our dataset was attributable to such an increase in sensitivity to rhythmic input accompanying the overall increase in activation level in response to preferred stimulus (**Fig. 5A**).

**Figure 5.**
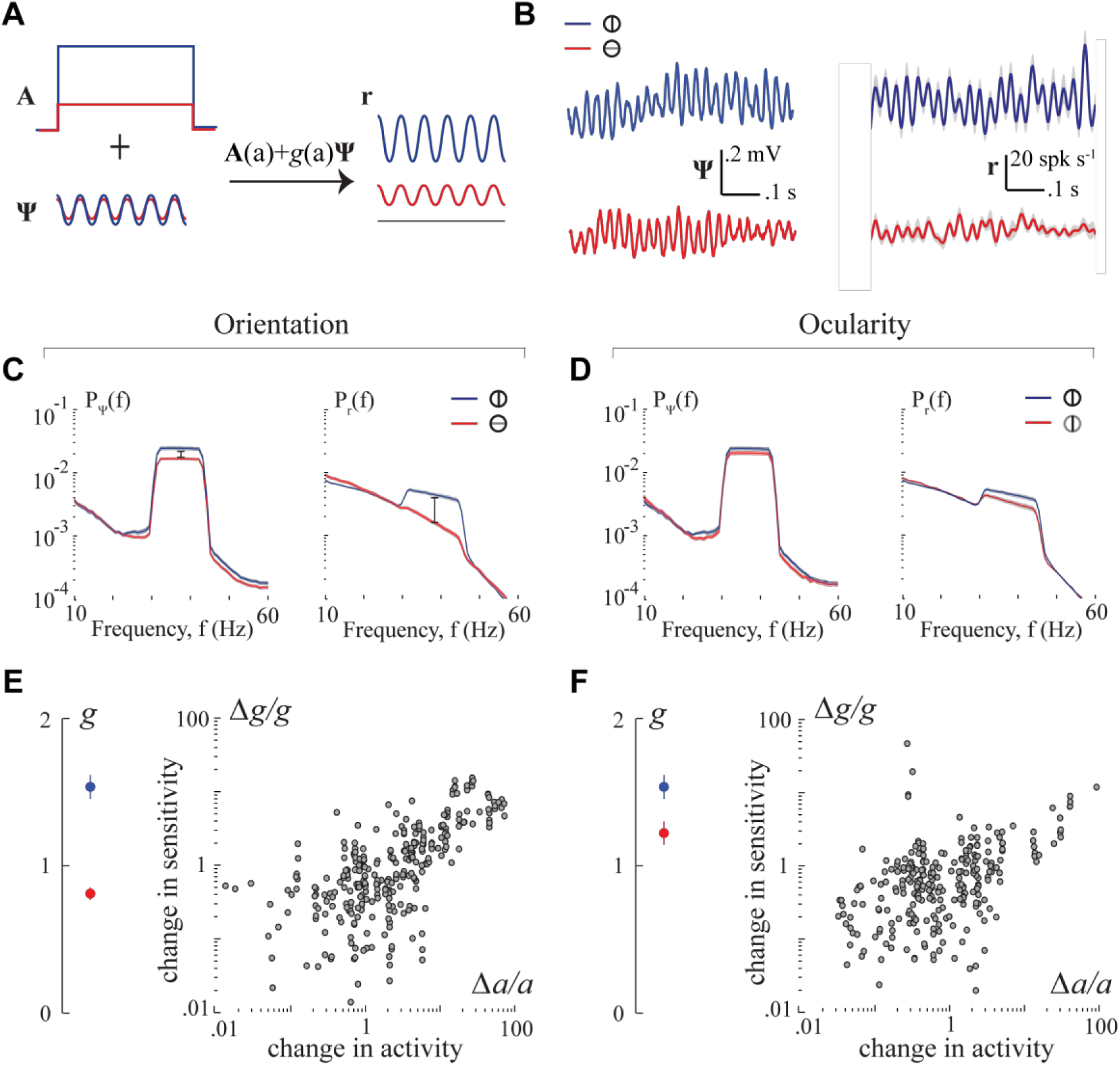
Modeling and analysis of oscillatory dynamics in spiking process. **(A)** Schematic of the model. Spiking process **r** is modelled as a sum of asynchronous excitation with mean *a,* and synchronous input **Ψ** amplified though a sensitivity parameter *g*. Red and blue traces correspond to non-preferred and preferred stimuli respectively. Stimulus-dependent increase in sensitivity can produce a large increase in synchronous firing even for modest changes in synchronous input. (**B**) A representative example. Left: LFP trace from each trial was time-shifted to match the phases of their gamma cycles before averaging to yield **Ψ** (see text). Right: Neuronal spike trains from the corresponding trials were shifted by the same amount as the LFP and averaged across trials to yield **r**. Mean values have been subtracted, and traces from the two conditions are vertically offset for visualisation. Shaded regions denote ± 1 standard deviation. (**C**) Power spectral densities of **Ψ** and **r**, averaged across the dataset for the pair of orientations. High firing rates in response to preferred stimuli resulted in a broadband increase in spectral content of **r**. Therefore for each condition, **P_r_** was normalised by the mean firing rate of the neuron before averaging to visualise changes in the gamma-band. Whereas stimulus-related modulation in gamma-band power of **Ψ** was modest, that of **r** was substantial. Shaded regions correspond to ±1 SEM. (**D**) Same as (C) for the pair of ocularity conditions. (**E**) Across the population of all spike-LFP pairs, mean sensitivity *g* to synchronous input was substantially greater in response to the preferred orientation (blue) than non-preferred (red). Error bars denote ±1 SEM. Fractional change in sensitivity was significantly correlated with change in activity (mean firing rate) as illustrated in the scatter plots. Each circle corresponds to one spike-LFP pair. (**F**) Same as (E) for the pair of ocularity conditions.

For each spike-LFP pair, we used the LFP signal as a proxy for **ψ** and inferred model parameters *a* and *g* that explained the spiking activity **r**. There were two challenges in this approach. First, while average synaptic input constitutes the major source of extracellular currents, LFP is also likely to contain traces of currents emerging from calcium spikes, action potentials, and spike after-potentials. Since the gamma rhythmic synaptic inputs are not precisely phase-locked with stimulus onset, we cannot use stimulus-locked trial-averaging to isolate the component reflecting rhythmic synaptic input. Second, spikes from individual trials are typically too sparse and noisy to allow reliable assessment of the underlying temporal dynamics. Once again due to the lack of phase consistency between rhythmic spiking activity and stimulus onset, trial-averaged firing rates typically fail to reveal the rhythmic process underlying spike generation (see Fig. 1C for example). We used the following technique to overcome both issues. We time-shifted our LFP traces on each trial by a small amount so that the peak gamma frequency in the LFP was phase-matched across trials. Spike trains from corresponding trials were shifted by the same amount to ensure that this procedure preserved the phase consistency of spike-trains relative to LFP i.e. spike-field coherence (**Supplementary Fig. 6**). **Figure 5B** shows the trial-averaged traces of the resulting time-shifted LFP (**ψ**) and spiking process (**r**) reconstructed above for a representative spike-LFP pair. In this example, it is clear that although the increase in amplitude of gamma oscillations in **ψ** was modest, there was a marked increase in the rhythmicity of **r** in response to preferred stimulus. This was observed across our dataset and can be noticed in the population averages of the power spectra of **ψ** and **r** (**Fig. 5C, D**).

If sensitivity *g* was independent of stimulus, the relative change in spectral content of **r** would be comparable to those in **ψ**. However, the fractional increase in gamma-band power was larger in **r** suggesting that there was a stimulus-dependent increase in sensitivity. To confirm this, we fit the activation parameter *a* and sensitivity *g* individually for each spike-LFP pair. Whereas *a* was trivially given by the mean of the spiking process **r**, sensitivity *g* was inferred by computing the fraction of temporal variability in spiking that was contributed by the rhythmic input (**Supplementary Fig. 7, Methods equation** (**5**)-(**6**)). As shown in **Figure 5E, F** (left panels), there was indeed a significant increase in sensitivity to rhythmic input in response to preferred stimulus (*p*<10^-5^; Wilcoxon rank-sum test). Moreover, changes in sensitivity were strongly correlated with the increase in neuronal activation *a* (Pearson’s correlation - orientation: *r*=0.59, *p*<10^-5^; ocularity: *r*=0.31, *p*<10^-5^) (**Fig. 5E, F** - right), and not to the strength of rhythmic input (orientation: p=0.78; ocularity: p=0.57) (**Supplementary Fig. 8**). Does the increase in sensitivity specifically underlie the increase in neuronal coherence? We tested this by partitioning the spike-LFP pairs into two groups, based on whether stimulus-dependent changes in neuronal coherence were congruent or incongruent with firing rates. Whereas preferred stimuli elicited a large increase in sensitivity in the congruent pairs, the effect on incongruent pairs was much smaller and barely approached significance (orientation: *p*=0.07, Wilcoxon rank-sum test, *n*=69 spike-LFP pairs; ocularity: *p*=0.09, *n*=100; **Supplementary Fig. 9**) confirming that decrease in coherence with firing rates in these pairs were attributable to the lack of increase in sensitivity to rhythmic input. Together, these results suggest that there is activity-dependent increase in sensitivity to rhythmic input, and that this specifically underlies the strong correlations between firing rate and neuronal coherence.

## Discussion

We examined concomitant changes in gamma-band neuronal coherence, firing rate, and strength of extracellular gamma rhythms in the primary visual cortex of awake rhesus macaques viewing monocularly presented grating stimuli. We found three key results that all point to an active mechanism that modulates the transfer of synchrony by neurons depending on their overall activation level.

First, neuronal gamma-band coherence quantified using spike-field coherence (SFC) increased with firing rate such that stimulus identity (both orientation and eye) was encoded in both quantities. Although neuronal coherence has previously been observed to increase with firing rate, some studies have found that rate and coherence encode different features of the stimulus leading to the view that the two codes carry complementary information^23–25^. In contrast, our result suggests that the two may also operate in tandem to increase information throughput about the same stimulus feature.

Second, neuronal coherence was more strongly correlated with firing rate than with the strength of gamma rhythms in the local field potential (LFP). This is surprising because gamma rhythms in the LFP are thought to reflect synchronous synaptic inputs^26,27^ that orchestrate rhythmic synchronization between neurons. On the other hand, SFC measures coherence in the spiking activity and largely reflects the amount of coherent output of the neurons. The fact that SFC was better predicted by firing rates suggests that the net coherence transferred by neurons depends more strongly on their overall activation level than on the strength of synchronous input. In one previous study, individual neurons in cortical slices were stimulated by injecting current steps of different means combined with noisy gamma oscillatory currents to study how the oscillations interact with the overall activation level^28^. The authors observed that the activation level had a significant impact on the timing of spikes elicited by gamma oscillatory input in a manner that increased the probability of coincidental spiking between neurons with similar firing rates, a phenomenon they called “rate-specific synchrony”. Although their work does not pertain to oscillatory neuronal synchronization and therefore distinct from ours, it supports the notion that activation level could have a significant impact on determining how gamma oscillatory input affects neuronal spiking.

Third, the spatial scale of gamma-band neuronal coherence was much smaller than extracellular gamma rhythms, but indistinguishable from that of the firing rate code. If coherent spiking were a consequence of passive integration of rhythmic synaptic inputs reflected in the LFP, it would share the same spatial scale as extracellular rhythms. The fact that the spatial correlation of neuronal coherence was instead comparable to that of firing rates reinforces the view that neuronal activation level must play a key role in the gating of rhythmic inputs. It follows that the local differences in activation level would decorrelate long-range coherent fluctuations in gamma rhythmic synaptic input to restrict the spatial scale of coherence in spiking activity. Although the spatial scale of LFP gamma rhythms in our measurements is consistent with earlier work, it has been shown that this scale is in fact variable and depends on the size of the stimulus^29,30^. While it is likely that the precise scale of extracellular rhythms in our dataset is specific to our choice of stimulus size, we believe this does not affect our interpretation. In fact, large spatial correlations in LFP rhythms induced by our stimulus helped capture the decoupling between the spatial scale of neuronal coherence and the LFP. We note that the pattern of results was qualitatively similar regardless of whether changes in responses were brought about by changes in orientation or ocularity. Given that the spatial scale of orientation and ocular columns in macaque V1 differ by at least an order of magnitude^31,32^, our findings here are likely to extend beyond the encoding of specific stimulus features. Together, our experimental results all point to some form of interaction between neuronal activation-level and gating of rhythmic inputs.

We tested this possibility by developing a novel statistical technique to fit a linear model that explicitly captured the dependence of rhythmic spiking on both the overall activation level as well as the strength of synchronous input through a sensitivity parameter. We found that stimulus-related increase in gamma rhythmicity of the spiking process could not be entirely accounted for by an increase in strength of rhythmic synaptic input but stemmed largely from an activation-dependent increase in neuronal sensitivity to rhythmic drive. Although our model is phenomenological, past experiments suggest that our findings may have mechanistic origins at the neuronal level. Intracellular recordings in vivo have demonstrated an activity-dependent increase in spike threshold that facilitates coincidence detection^33–37^. Consistent with this, biophysical modeling studies and slice recordings show that increase in Poisson-like synaptic background activity increases the sensitivity of pyramidal neurons to temporally structured synaptic input^38–40^ by effectively increasing voltage threshold via *M* currents^41,42^. It is possible that such dynamic changes in synaptic integration properties may also underlie the ability of neurons to adapt their sensitivity to gamma rhythms reported here. In fact, recently Perrenoud et al used intracellular recordings to demonstrate that increased phase-locking of pyramidal neurons to the gamma cycle is facilitated by an overall increase in the average membrane potential in response to visual stimulation^43^. Alternatively, the observed changes in response characteristics of neurons and the associated gamma-band synchronization could be a reflection of network-level computations such as divisive normalization^44^. If this is the case, the normalization pool is likely to be confined to within an orientation column for otherwise we would not have observed orientation-dependent changes in neuronal sensitivity. Further experiments will be necessary to identify whether the observed phenomenon is dominated by a cellular or network mechanisms.

There are some fundamental issues in relation to past findings which merit further scrutiny. First, increase in neuronal gamma-band synchronization with firing rate has been reported previously^5,6,12,45,46^. Theoretical studies have shown that this dependence is expected for a broad class of statistical models^47,48^. Mechanistically, such dependence could stem from a simple threshold nonlinearity in the neurons^49^. However, simulations of model neurons with a fixed threshold substantially underestimated the magnitude of rate-dependent increase in SFC observed in our data (**Supplementary Fig. 10-11**). Instead, our data was better explained by a model in which neuronal threshold increased with the mean activity of the neurons (**Supplementary Fig. 12**). This dynamic threshold model is consistent with several past experimental results, and supports the idea that the mechanism mediating neuronal coherence is sensitive to neuronal activation. Second, since synaptic currents constitute a major source of fluctuations in the LFP^18–20^, we used gamma oscillatory power in the LFP as a proxy for the strength of rhythmic synaptic input to neurons. Although the precise magnitude of rhythmic input to individual neurons likely differs from that estimated using the LFP^20^, our conclusions are valid insofar as the relative changes in LFP oscillations are correlated with those of oscillatory input to single neurons across stimulus conditions. It is difficult to test this precisely without intracellular recordings. However it has previously been shown that the average LFP waveform around the time of a neuron’s action potential (spike-triggered LFP) reflects the correlation between the LFP and that neuron’s synaptic inputs^22^. The STAs in our dataset had particularly large gamma-band powers and their amplitudes were not significantly modulated by stimulus, so we believe gamma activity in the LFP provided a robust readout of the strength of gamma rhythmic synaptic inputs. Finally, neuronal coherence in the gamma band is known to vary independently of firing rates in some cases^11,24,50,51^ suggesting that top-down factors such as attention may also potentially enhance coherence by altering neuronal sensitivity in a similar manner. Moreover, factors such as noise-level and size of stimulus have been shown to alter the spatial scale of LFP leading to more spatially tuned gamma activity^29,30,52^. In such cases, changes in coherence can also come about more directly from an increase or decrease in rhythmic synaptic input to neurons regardless of their activation-level. Quantitative analyses using an approach similar to ours will be necessary to clarify the mechanistic origins of rate-independent changes in coherence.

The potential implications of gating coherence based on activation-level are at least twofold. First as we have shown, this would lead to a similarity in the spatial resolution of firing rate and gamma synchrony codes. It has been argued that the brain can decode temporally multiplexed codes using known mechanisms^4^ such as synaptic depression and facilitation that endow neurons with multiple synaptic timescales^53,54^. Performing spatial de-multiplexing in addition to the above would not be as easy because it would require differential spatial pooling depending on the timescale. Moreover it is unclear if such algorithms might be supported by the brain’s neural hardware. The proposed mechanism obviates the need to deal with such complexities by matching the spatial scales of the two codes. Second, current theories suggest that gamma synchrony may be involved in the encoding of sensory information^55–57^, regulating information flow between brain areas^58^^-^^60^, and facilitating synaptic plasticity ^6,61,62^ Supporting such a diverse range of functions would require synchrony to be robust to the precise level of neuronal activity. Adaptive gating of coherence based on activation-level would also help achieve this by preserving the transfer of synchrony in the face of elevated asynchronous background activity.

## Methods

### M1 Electrophysiological recordings

Two adult male rhesus monkeys (*Macaca mulatta*) D98 and F03 weighing 16 kg and 11 kg respectively, took part in the experiments. Cranial headposts and form-specific chambers were surgically implanted. Recording chambers were positioned stereotactically over the operculum in area V1 in both hemispheres of D98 and right hemisphere of F03 with the aid of high-resolution anatomical scans. Skull parameters extracted from these scans were used for designing the headpost and the recording chambers to fit the skull surface. A more detailed description of these methods can be found elsewhere ^63–65^. A custom-built array of tetrodes ^66^ was chronically implanted in area V1 inside the recording chamber implanted in the left hemisphere of the monkey D98. The tetrodes were at least 200μm apart ^67^ Recordings were also carried out non-chronically from the right hemisphere of both monkeys. In these sessions, one to four manually adjustable microdrives (Crist Instrument Co.) were inserted into a custom-built grid and the activity was recorded using tetrodes. The experimental and surgical procedures were performed with great care and in full compliance with the German Law for the Protection of Animals, the European Community guidelines for the care and use of laboratory animals (EUVS 86/609/EEC), and the recommendations of Weatherall report ^68^. The regional authorities (Regierungspräsidium Tuebingen) approved our experimental protocol application and the institutional representatives for animal protection supervised all procedures.

The raw voltage signal was passed through an analog bandpass filter (1-475 Hz), sampled at ~1990.7 Hz, digitized (12 bits) and stored as the LFP signal. Multiunit spikes were identified by passing the raw signal through a separate analog bandpass filter (600 Hz-6 KHz), followed by sampling (32 KHz), digitization (12 bits) and detecting the times at which the signal crosses a predefined threshold (25μV). Following each threshold crossing, a segment of 32 samples (1ms) was extracted from all four channels of the tetrode and these waveforms were stored for offline clustering. Single-unit spikes were then isolated from multiunit activity by a custom-built clustering software ^66^ that uses features extracted from the stored multiunit spike waveforms.

### M2 Visual stimuli

A dedicated graphics workstation (TDZ 2000; Intergraph Systems) running an OpenGL-based program was used for rendering visual stimuli, while the behavioral aspects (e.g. juice reward, trial abortion) were controlled using the QNX real-time operating system (QNX Software Systems Ltd). The display system comprised of a custom-made mirror stereoscope with an LCD monitor (resolution of 1024x768; refresh rate of 60 Hz) on each side as shown in **Fig. 1B**, and allowed for dichoptic presentation of stimuli.

Each session began with a calibration procedure ^63^ to ensure that the monkeys could correctly overlay (fuse) the central fixation markers (0.2°) on the two displays. The following procedure was then carried out to determine the position and orientations of the stimuli to be used in the experiments. A grating of arbitrary orientation was presented binocularly (i.e., shown to both eyes) at a parafoveal location while the monkey fixated on the central marker. The location, size and orientation of the grating are then systematically changed until the location of the receptive field and the orientation preference of the multiunit response could be estimated. Such online estimation was made possible by playing the multiunit activity through a sound amplifier (Grass Technologies). The pair of orthogonal orientations (*θ_A_* and *θ_B_*) that elicited maximal differential multiunit response were identified and used in the experiment.

Each trial of the experiment began with the monkey fixating on a central marker (0.2°). After maintaining fixation for 300ms, a static sine-wave grating stimulus (diameter of 1-2°; spatial frequency of 3-5 cycles/deg; contrast 70%) of two possible orientations was displayed monocularly to one of the eyes for a period of one second (**Fig. 1B**). The animal was required to maintain fixation within a circular window with a radius 0.5° from the center of the marker throughout the duration of the trial. At the end of each successful trial, a drop of apple juice was delivered as a reward. A failure resulted in abortion of the trial without reward. Depending on the orientation of the grating and eye of presentation, each trial belonged to one of four stimulus conditions (**Fig. 1B**). A typical recording session included 200 trials of each condition. Throughout this paper, the term ‘preferred’ orientation (eye) is used to refer to the orientation (eye) that elicits higher firing rate for a given singleunit. The complementary condition is dubbed ‘nonpreferred’.

### M3 Data Analysis

Singleunits were first tested for visual responsiveness by comparing stimulus-evoked firing rates to baseline. Only neurons that exhibited a significant increase in their firing rates in response to visual stimulus (*p*<0.05; Wilcoxon rank-sum test) were considered for further analyses. Unless otherwise specified, all time-domain and spectral estimates were based on responses recorded between 400-1000ms following stimulus onset when signals were relatively more stationary.

#### Spike-field coherence (SFC)

To measure the extent of rhythmic synchronization between LFP and spike trains at all frequencies, we estimated spike-field coherence (SFC) defined as the squared magnitude of the cross-spectrum divided by the product of the auto-spectra ^69^:

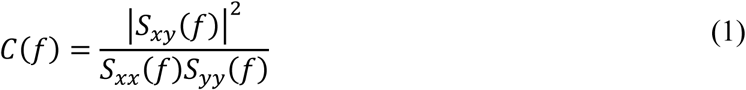

where *C*(*f*) denotes the spike-field coherence at frequency *f, S_xy_*(*f*) denotes the cross-spectral density function between spike train *x* and LFP signal *y, S_xx_*(*f*) and *S_yy_*(*f*) are the respective autospectra. All spectral estimates were carried out using multi-taper method with *K* = 7 orthogonal Slepian tapers **w***_k_* to yield spectral smoothing of approximately ±4 Hz at a frequency resolution of ~1 Hz. This involved multiplying each data sequence **x***_n_* (**y***_n_* for LFP) with the different tapers to obtain *K* independent spectral estimates and then averaging them:

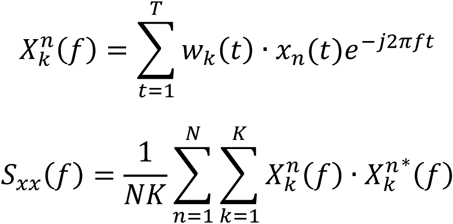

where 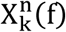 denotes the *k^th^* Slepian-tapered Fourier transform of **x***_n_*, 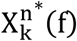 its complex conjugate, *N* is the total number of trials, and *T* is the duration of the signals *x* and *y.*

#### Significance test for SFC

To test the statistical significance of SFC for each spike-LFP pair at every frequency, we obtained multiple estimates of SFCs by shuffling the order of the trials of spike trains thereby destroying the phase relationship between spike trains and LFPs. At any given frequency, the estimated value of SFC before shuffling was deemed to be significant if the probability of drawing it from the distribution of shuffled SFC estimates was less than 0.01. Only those pairs which showed significant SFC at 8 consecutive bins (~8Hz) in the frequency range between 30-45 Hz in at least one of the stimulus conditions were considered for further analyses. We assessed the significance of difference in SFC across the two pairs of conditions by a shuffling procedure similar to the one described above. Trials from the two conditions to be tested were pooled together. Half the trials were then dubbed to be from one condition, while the other half were labeled to be from the other condition. SFCs were then estimated for the two conditions and the differences in SFCs were computed. This procedure was repeated several times and the true difference was compared against the distribution of ‘fake’ differences (*p*<0.01). A similar technique was used to assess significance of modulations in gamma-band LFP power. Statistical significance of firing rate changes were tested using two-sided Wilcoxon rank-sum test (*p*=0.05) by comparing spike counts across trials for the relevant stimulus conditions.

#### Modulation Indices

Orientation and ocularity preferences were first determined for each neuron by estimating rate-modulation indices (*M_R_*) derived from the average firing rates (*R*) elicited by the pair of stimuli:

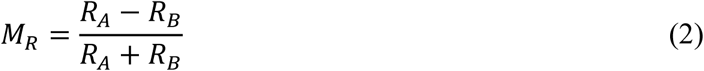

Here *A* and *B* are used as placeholders to denote the pair of conditions that correspond to the presentation of, either the pair of orthogonal gratings to the neuron’s preferred eye (orientation preference), or the preferred orientation to the right and left eye (ocularity preference). Similar definitions were used for quantifying modulations in the gamma-band LFP power (*M_L_*) and gamma-band spike-field coherence (*M_c_*). To explicitly test whether modulations in firing rate (*M_R_*) or LFP power (*M_L_*) better predicted modulations in neuronal synchrony (*M_c_*), we performed multiple linear regression *M_c_* = *β_R_M_R_* + *β_L_M_L_* + *β*_0_ to determine coefficients *β_R_* and *β_L_* on predictors *M_R_* and *M_L_* respectively, according to:

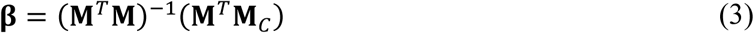

where **β** = (*β_R_ β_L_ β*_0_)*^T^*, and **M** = (**M***_R_* **M***_L_* **1**) where **M***_R_* and **M***_L_* denote vectors of modulation indices across the population. To compare the quality of predictions given by *M_R_* and *M*_l_, we regressed *M_c_* against *M_R_* and *M_L_* separately. The mean slopes of both regressions were used individually to generate predictions 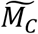. For each spike-LFP pair, normalised residuals were then estimated by computing the squared deviation of the prediction from the experimental measurement *M_c_*, normalized by the variance of the measurement: 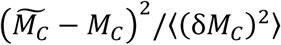.

#### Pseudo spike-field coherence

In addition to assessing modulation of firing rates and SFC across conditions, we directly tested whether firing rates exhibited correlated variability with SFCs across trials within a given condition. This was done by calculating the Spearman correlation coefficient (*ρ*) between spike count across trials and single-trial coherence estimates. Coherence estimates for individual trials were obtained through the following procedure ^5^. For any given trial, the z-transform of the SFC estimated by leaving out that trial, was subtracted from the original z-transformed SFC estimate after weighting each term with the number of trials used in the estimate:

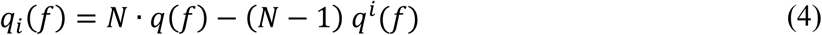

where *q_i_*(*f*) denotes pseudo-coherence of the *i^th^* trial, *q*(*f*) is the coherence across all *N* trials, and *q^i^*(*f*) is coherence estimated by leaving the *i^th^* trial out. Here 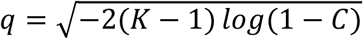 denotes the z-transformed value of the estimated coherence *C* with *K* = 7 tapers.

#### Spike-triggered LFP

For each neuron, spike-triggered average (STA) of the LFP was estimated first by averaging 300ms segments around the time of each spike emitted by that neuron. The result was divided by the standard deviation of the LFP to obtain normalised STA^22^. The peak amplitude of the normalised STA was taken as a measure of the correlation between LFP and neuronal membrane potential. To determine whether LFP was robustly correlated with the membrane potential across all stimulus conditions, normalised STAs were computed separately for each stimulus condition and their peak amplitudes were compared.

#### Correction procedures

To ensure that our test statistics are not affected by differences in bias and variance across conditions, we took the following measures to avoid potential confounds in SFC estimation. Each condition included the same number of trials and the lengths of data segments in all trials were identical across conditions. For each single-unit, SFCs were calculated between its spike trains and concurrently recorded LFPs from all tetrodes. Consequently ~70% of SFCs (275/400 pairs) were obtained from spikes and LFPs belonging to different electrodes. Furthermore, when using spikes and LFP from the same electrode, a 4ms segment of LFP was removed around the time of each spike and those data points were replaced using cubic spline interpolation. This procedure resulted in a significant reduction of SFC, but only in the frequency range above 100 Hz (**Supplementary Fig. 13**) implying that gamma-band SFC was not affected by spurious correlations between spikes and LFP. Therefore, we retained the spike-LFP pairs for which spikes and LFP were recorded on the same electrode in order to gain statistical power.

### M4 Neuronal model

Rhythmic spiking process *r*(*t*) was modeled as a sum of asynchronous excitation *A*(*t*) with mean *a* and rhythmic input *ψ*(*t*) as described in the results. The net excitability due to *ψ*(*t*) is assumed to be zero, so the mean activity is given by 〈*r*(*t*)〉 = 〈*A*(*t*)〉 = *a*. Variability on the other hand, is inherited from both inputs according to:

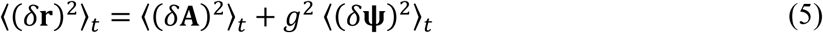

where functions of time are denoted using boldface letters for convenience, *g* is the sensitivity to synchronous input, and subscript 〈·〉*_t_* denotes expectation over time. To estimate 〈(*δ***r**)^2^〉*_t_*, we first time-shifted the spike train of each trial *i* by an amount *δt_i_* so that gamma phases of the simultaneously recorded LFP traces were aligned across trials, and then computed the temporal variability of the trace obtained by averaging the resulting spike trains across trials (**Supplementary Fig. 7A** – top right). We isolated the component of variability due to rhythmic input by subtracting the asynchronous component 〈(*δ***A**)^2^〉*_t_* from 〈(*δ***r**)^2^*〉_t_*. The asynchronous component was estimated by a procedure in which elements of {*δt_i_*} were shuffled before shifting the spike trains. This shuffling procedure essentially randomizes the phases of gamma cycles of different trials, so that any temporal variability in the resulting trial-averaged firing rates will be solely due to asynchronous dynamics of the spiking process (**Supplementary Fig. 7A** – bottom left). The shuffling procedure was carried out several times and the mean of the resulting distribution of variabilities 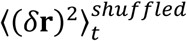 was used as an estimate of 〈(*δ***A**)^2^〉*_t_.* Neuronal sensitivity *g* was then estimated according to:

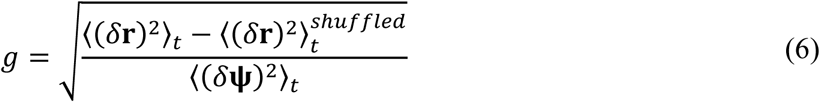

where 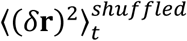 denotes variability estimated through the shuffling procedure.

## Acknowledgements

We would like to thank Drs. Jacob Macke, Philipp Berens, Esther Florin, and David Omer for comments on previous versions of the manuscript. This work was supported by the Max-Planck Society and the German Federal Ministry of Education and Research (BMBF; FKZ: 01GQ1002).

